# Virtual Screening for the Discovery of Novel Urease Inhibitors Targeting Rumen Bacterial Urease

**DOI:** 10.1101/2022.08.16.504210

**Authors:** Shengguo Zhao, Larisa A. Ilina

**Affiliations:** Institute of Animal Sciences, Chinese Academy of Agricultural Sciences, Beijing, China; St. Petersburg State Agrarian University, 196601 Pushkin, Russia

**Keywords:** *Ruminococcus albus* 8, urease inhibitors, virtual screening, molecular docking, molecular dynamics

## Abstract

The inhibition of urea hydrolysis by rumen bacterial urease presents a promising strategy to enhance dietary nitrogen utilization efficiency in animal production while mitigating environmental nitrogen pollution. In this study, we employed comparative modeling using Modeller to determine the three-dimensional structure of urease from the ruminal bacterium Ruminococcus albus 8 (RaUrease), based on the crystal structure of Helicobacter pylori urease (HpUrease, PDB ID 1E9Z). Through molecular dynamics simulation conducted for 20 ns using the AMBER force field within the Gromacs system, we generated a reliable RaUrease structure and subsequently evaluated the stereochemical quality of the protein model. Virtual screening identified 12 commercially available compounds as potential urease inhibitors, based on their binding energies (<-9.8 kcal mol-1). Characterization of the RaUrease-binding site for the compound exhibiting the highest predicted inhibitory activity revealed that van der Waals interactions play a more significant role than hydrogen bonding in rumen bacterial urease binding. These findings provide valuable insights for the design of novel urease inhibitors tailored for ruminants, offering potential solutions to reduce nitrogen pollution associated with livestock production.

## 1. Introduction

Urease (Urea Amidohydrolase, EC 3.5.1.5) is a thiol-rich, nickel-dependent metalloenzyme that catalyzes the hydrolysis of urea into ammonia and carbamate. The catalytic activity of urease is dependent on the presence of nickel ions (Ni^2+^) and sulfhydryl groups, particularly the multiple cysteinyl residues located in the enzyme’s active site. Structural studies of *Helicobacter pylori* urease (HPU) have revealed a dinuclear nickel active site, where a carbamylated lysine residue bridges the two deeply buried metal atoms. The active site is surrounded by hydrophilic amino acids and features a highly flexible flap that undergoes an induced fit upon substrate binding. It has been demonstrated that the catalytic activity of urease is strongly dependent on its multiple cysteinyl residues bearing sulfhydryl groups, particularly those located on the mobile flap that closes over the active site.

Urea has been widely used as a nitrogen supplement in ruminant rations for over a century to increase ruminal degradable nitrogen and reduce feed costs [1]. However, the use of urea in ruminant diets must be approached cautiously. Excessive urea hydrolysis, driven by bacterial urease in the rumen, can surpass the capacity of ammonia assimilation, leading to inefficient nitrogen utilization, blood ammonia toxicity in animals, and environmental nitrogen pollution [2, 3]. To address these issues, various urease inhibitors, such as acetohydroxamic acid (AHA), phenyl phosphorodiamidate (PPDA), and N-(n-butyl)thiophosphoric triamide (NBPT), have been investigated as dietary adjuncts in animal husbandry over the past decades.

Unfortunately, many of these inhibitors provide only short-term control, as their efficacy diminishes over time due to microbial adaptation [4–7]. Consequently, there is a pressing need to identify novel, safe, and effective urease inhibitors with long-term activity for use in dairy production.

The discovery and validation of novel urease inhibitors for ruminants typically require long-term livestock experiments, which are both costly and time-consuming [8]. Therefore, it is crucial to narrow down potential candidates before conducting extensive animal trials. Virtual screening offers a complementary approach to identify novel inhibitors by rapidly and reliably docking commercially available compounds into the active site of the target protein [9]. This method significantly reduces the number of candidate compounds that need to be tested experimentally [10]. For virtual screening to be effective, a three-dimensional (3D) structural model of the target protein—in this case, rumen bacterial urease—is highly desirable.

However, isolating and identifying rumen ureolytic bacteria is challenging, and our understanding of the structural and functional properties of ureases from ruminal bacteria remains limited. Among the rumen bacterial community, *Ruminococcus albus* 8 is a key bacterium involved in cellulose decomposition and microbial protein synthesis. Bioinformatics analysis of its genomic sequence suggests that it can synthesize urease [11]; however, the urease from this bacterium has not been characterized, and its crystal structure is not available in the Protein Data Bank. Previous studies have shown that ureases from different bacteria exhibit high amino acid sequence homology, with their active sites located in the α subunits [12]. In the absence of an experimentally determined structure, comparative homology modeling provides a valuable tool to generate a 3D model of a protein based on the structure of a related protein with known conformation [13]. Thus, homology modeling can be applied to predict the 3D structure of urease from rumen ureolytic bacteria.

In this study, we employed homology modeling combined with molecular dynamics simulations to generate the 3D structure of urease from *R. albus* 8 (RaUrease). Molecular docking was performed to predict the binding site of RaUrease using a known inhibitor as a positive control. Subsequently, virtual screening was conducted to identify novel and potential inhibitors of rumen bacterial urease. The results of this study are expected to narrow down candidate compounds for further validation through long-term animal experiments.

## 2. Results and Discussion

### 2.1 Sequence alignments and molecular modeling

RaUrease consists of 570 amino acids (Figure 1). When compared to the template sequence of HpUrease (568 amino acids), the two proteins exhibit a sequence similarity of 69%. Comparative models are considered highly accurate when the sequence identity to their templates exceeds 50%, as this typically corresponds to a root mean square deviation (RMSD) of approximately 1 Å for the main chain atoms. Most deviations in such models arise from side chain packing and minor shifts or distortions in the core regions of the main chain [14]. Therefore, the 69% sequence similarity observed in this study indicates a high level of homology, supporting the construction of a reliable structural model. The predicted three-dimensional (3D) structure of RaUrease was generated using Modeller 9.16, as illustrated in Figure 2(a).

**Figure 1.**
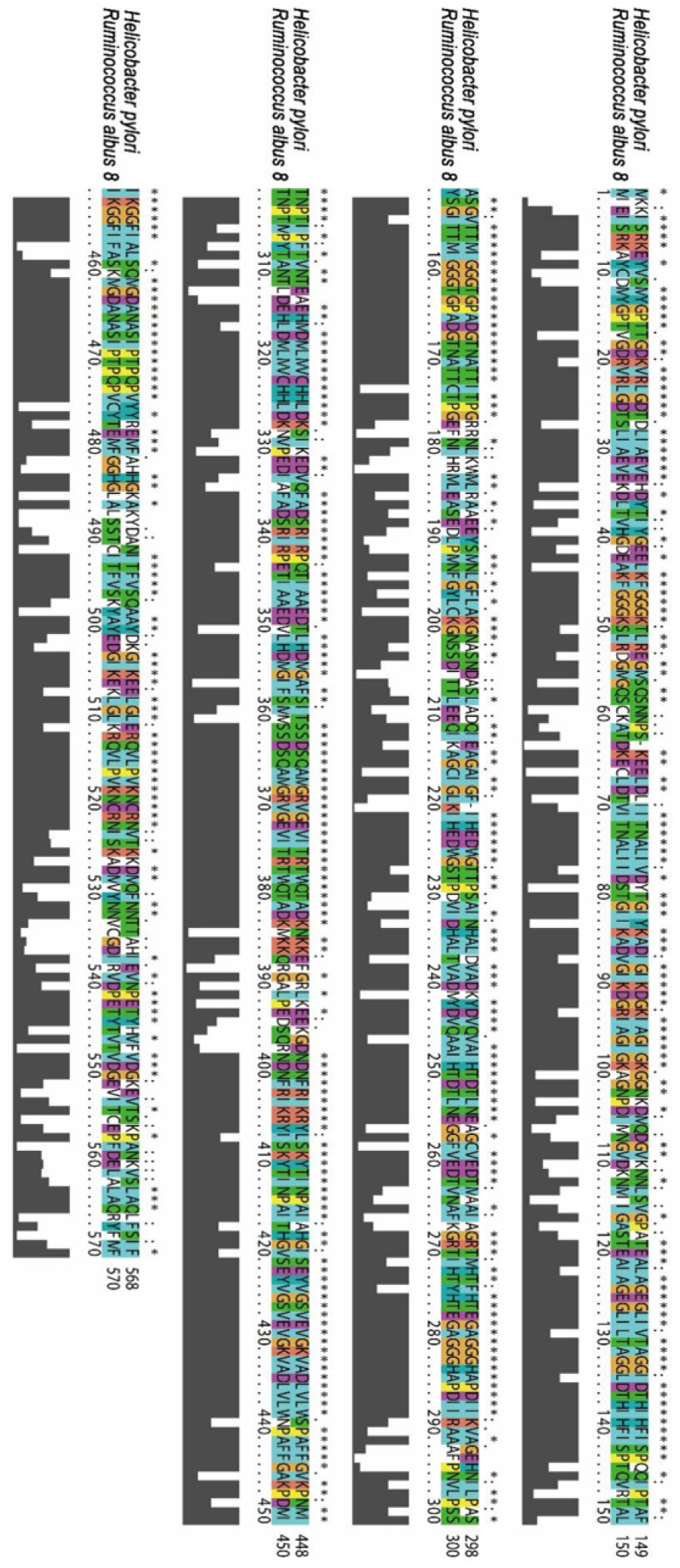
Sequence alignment between ureases from *Ruminococcus albus* 8 and *Helicobacter pylori*. The sequence identity at amino acid level was 69%.

**Figure 2.**
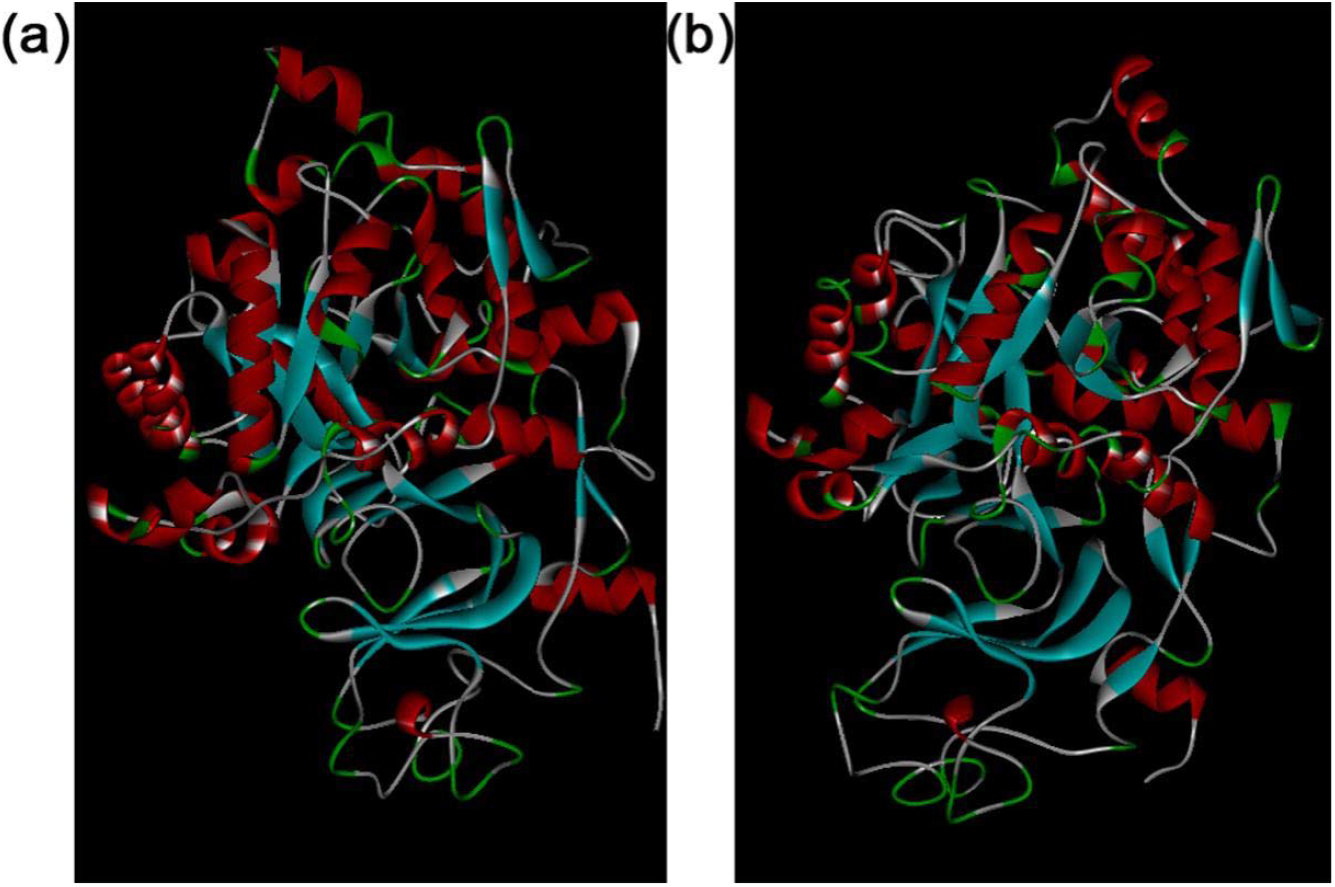
The predicted three dimensional (3D) model structure of *Ruminococcus albus* 8 urease was obtained using Modeller 9.16. The picture on the left (a) is the 3D structure established before molecular dynamics simulation, and the one on the right (b) is the structure after molecular dynamics simulation.

### 2.2 Optimization and validation of the homology model

In the molecular dynamics simulation study, conformational changes in the model structure of RaUrease were analyzed for a simulation period of 20 ns, and the predicted result is shown in Figure 2 (b). Ramachandran plot calculations were performed to evaluate the stereochemical quality of the final refined model using the initial model structure of RaUrease as the reference structure. The detailed residue to residue stereochemical quality of the protein structure was assessed using the Molprobity program, and the results are shown in Figure 3 and Table 1.

**Table 1.** Validation of the three dimensional structures of model and template ureases.

| Validation | RaUrease (0 ns) | RaUrease (20 ns) | HpUrease |
| --- | --- | --- | --- |
| Favored regions | 89.4% (508/568) | 88.2% (500/567) | 75.5% (426/564) |
| Allowed regions | 97.5% (554/568) | 98.6% (559/567) | 92.7% (523/564) |
| Ramachandran outliers | 2.46% (14/568) | 1.41% (8/567) | 7.27% (41/564) |
| Verify_3D | 86.14% | 89.28% | 86.62% |
| ERRAT | 82.740 | 90.556 | 85.536 |

The three dimensional structure of model urease from *Ruminococcus albus* 8 (RaUrease) was resolved (RaUrease, 0 ns) using the structure of urease from *Helicobacter pylori* (HpUrease) as template. The model structure was optimized using molecular dynamics simulation with a 20 ns procedure (RaUrease, 20 ns).

**Figure 3.**
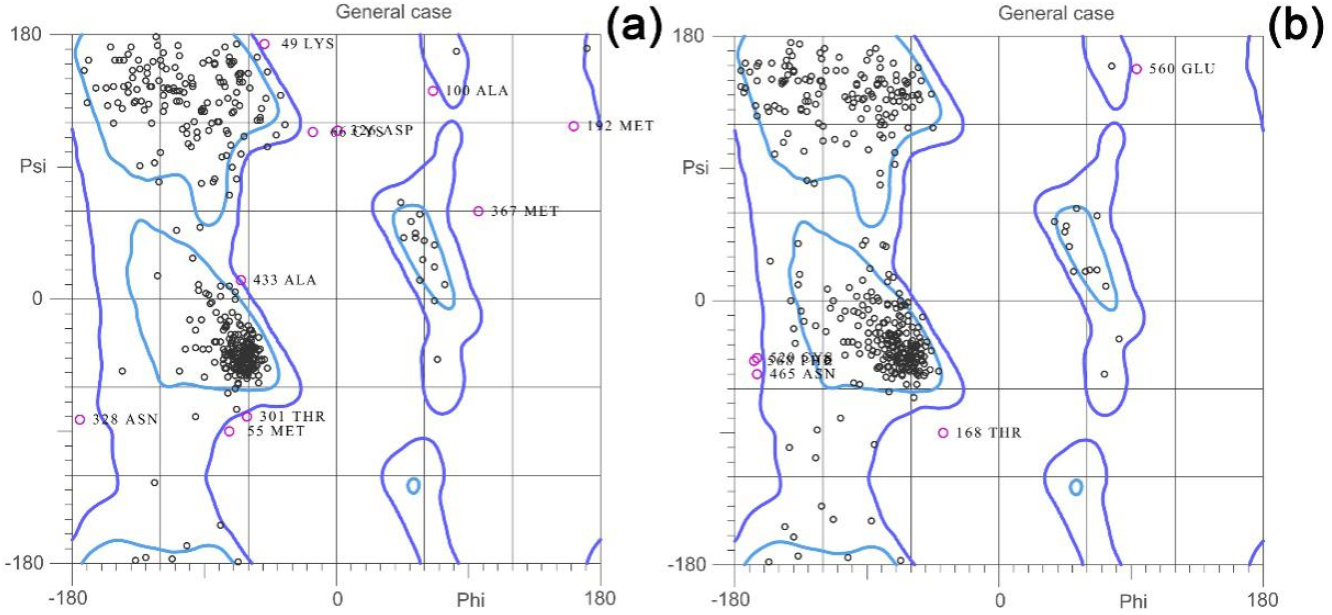
Ramachandran plots of *Ruminococcus albus* 8 urease initial (a) and final (b) three dimensional (3D) structures. There were 14 outliers (VAL17, LYS49, MET55, CYS66, ALA100, MET192, ILE287, THR301, ASN302, ASP326, ASN328, MET367, VaL427, ALA433) in the initial 3D structure of *R. albus* 8 urease, and eight outliers (ILE2, THR168, ASN302, PRO303, ASN465, CYS520, GLU560, PHE668) were still present after 20 ns of molecular dynamics simulation.

Before the optimization, the percentages of residues located in the favored, allowed, and disallowed regions were 89.4% (508/568), 97.5% (508/568), and 2.46% (508/568), respectively. After the refinement of the model protein, the percentages of residues in the favored, allowed, and disallowed regions were 88.2% (500/567), 98.6% (500/567), and 1.44% (500/567), respectively. There were 14 outliers (VAL17, LYS49, MET55, CYS66, ALA100, MET192, ILE287, THR301, ASN302, ASP326, ASN328, MET367, VaL427, and ALA433) in the initial 3D structure of RaUrease. Eight outliers (ILE2, THR168, ASN302, PRO303, ASN465, CYS520, GLU560, and PHE668) were still present after 20 ns of molecular dynamics simulation. The ERRAT scores changed from 82.740 to 90.556. The percentage of residues with average three dimensional-one dimensional (3D-1D) scores ranged from 86.14% to 89.28%, according to the results of Verify-3D. Generally, after the optimization, quality factors increased and error values decreased by satisfying the special restraints.

Ramachandran plot calculations were performed on the structure of the template urease from *H. pylori*, which was used to evaluate the reliability of the final refined model structure of RaUrease (Table 1). The percentages of residues for *H. pylori* urease located in the favored, allowed, and disallowed regions were 75.5% (426/564), 92.7% (426/564), and 7.27% (426/564), respectively. The ERRAT score was 85.536, and 86.62% of residues in HpUrease had average 3D-1D scores. The results showed that the geometric quality of the backbone conformation, residue interactions and contacts, and energy profiles of the final model urease were similar to those of the template protein, which suggests that a reasonable homology model of RaUrease was established. The verified model can be used for further investigations, such as protein-inhibitor interactions and virtual screening.

### 2.3 Identification of the binding region in the model urease by molecular docking

The ideal inhibitors are compounds that can fit well within the binding pocket. AHA is a simple and stable organic compound that has been used as a urease inhibitor in ruminant production in China since 1998 (No. 27 Letter on husbandry authorized by the Ministry of Agriculture of China). This representative inhibitor was docked to RaUrease using AutoDock Vina for the identification of the binding region in this model urease.

The RaUrease residues involved in the AHA-binding site are shown in Figure 4. AHA was surrounded by ALA390, LEU391, ASP394, SER395, GLN396, ASN398, ASP399, ASN400, PHE401, and ARG402. There were five hydrogen bonds formed between AHA and RaUrease. Among them, ASN398 formed two hydrogen bonds, and LEU391, PHE401, and ASP394 formed one each. The results imply that hydrogen bonds play an important role in AHA binding. The docking score for AHA was -4.5 kcal mol^-1^, which was used as the reference score for further virtual screening. The AHA-binding region on RaUrease was used as one of the active sites for further virtual screening. To identify additional potential inhibitors, the grid size for AutoDock Vina in the next molecular docking was set to cover the entire model enzyme structure.

**Figure 4.**
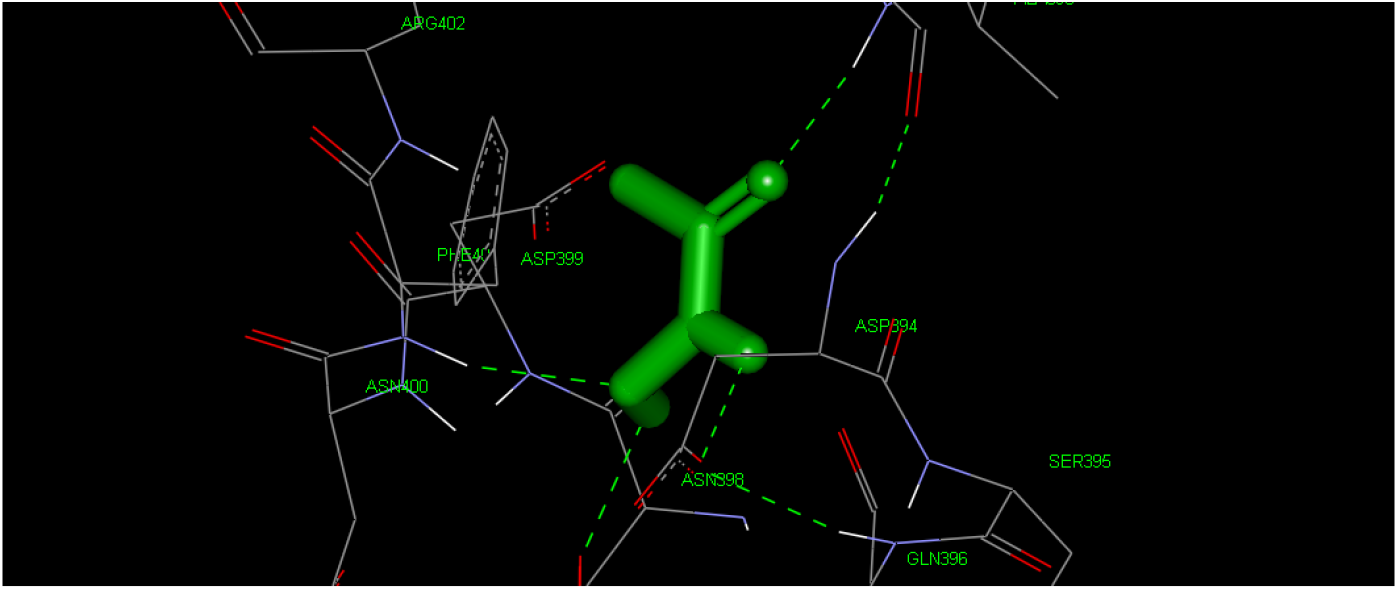
Predicted binding model of acetohydroxamic acid (AHA) to the model structure of *Ruminococcus albus* 8 urease. AHA was surrounded by ALA390, LEU391, ASP394, SER395, GLN396, ASN398, ASP399, ASN400, PHE401, and ARG402. There were five hydrogen bonds formed between AHA and the target protein.

### 2.4 Selection of potential urease inhibitors and docking of the highest affinity inhibitor to RaUrease

The high-throughput virtual screening yielded 22 candidate compounds with free binding energies of <-9.8 kcal mol^-1^. The 12 top hits are listed in Table 2. In comparison with the AHA score (−4.5 kcal mol^-1^), these screened candidates with better docking scores should have stronger interactions and better inhibitory efficiency. Some of the compounds among the 12 candidates were enantiomers, including the most efficient inhibitors, ZINC67911797 (−10.6 kcal mol^-1^) and ZINC67911804 (−10.1 kcal mol^-1^). To compare the two compounds, the binding patterns of ZINC67911797 and ZINC67911804 to the model RaUrease were determined and are shown in Figure 5.

**Table 2.**
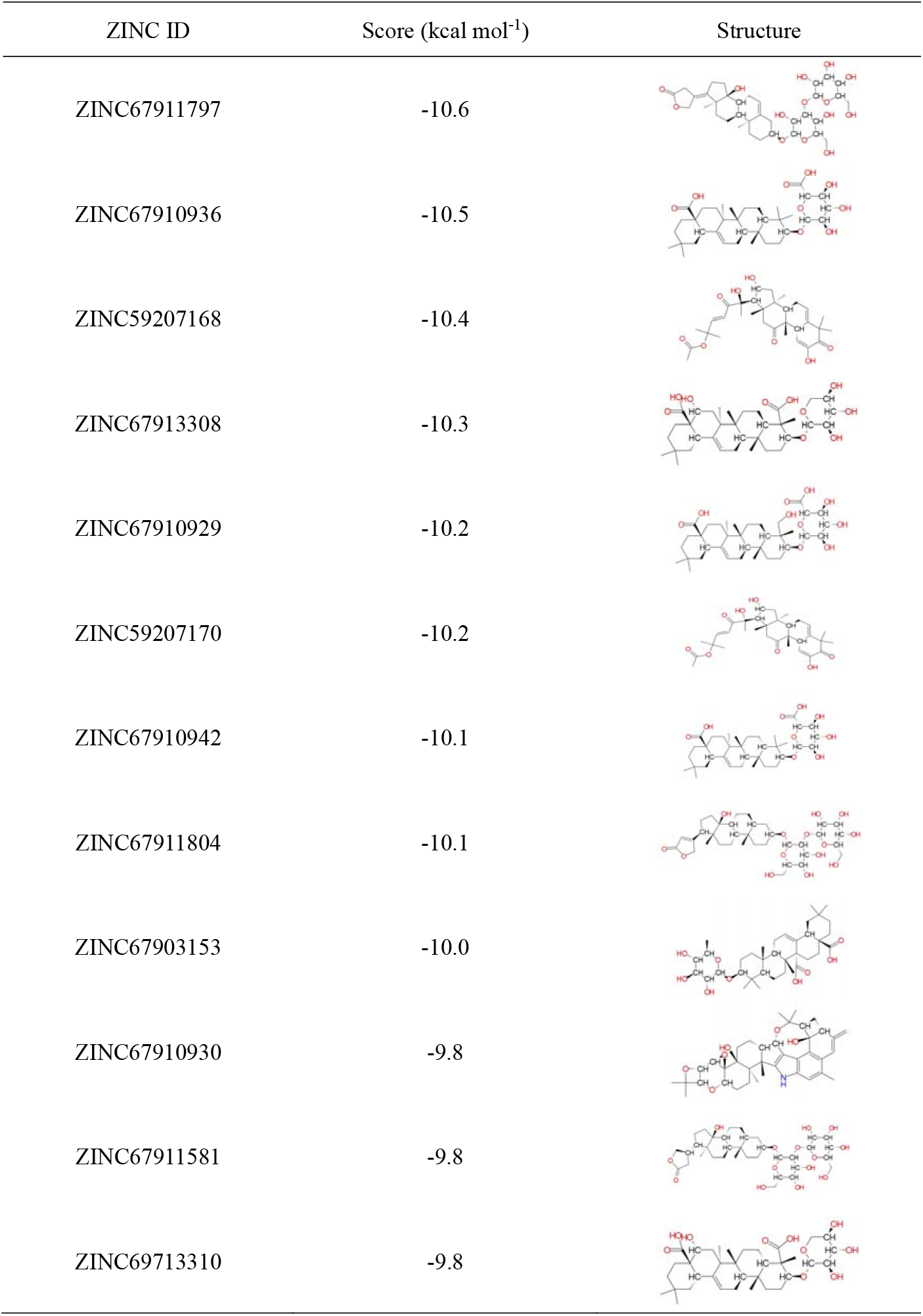
Free binding energy among novel urease inhibitors and the *Ruminococcus albus* 8 urease.

| ZINC ID | Score (kcal mol<sup>-1</sup>) | Structure |
| --- | --- | --- |
| ZINC67911797 | -10.6 |  |
| ZINC67910936 | -10.5 |  |
| ZINC59207168 | -10.4 |  |
| ZINC67913308 | -10.3 |  |
| ZINC67910929 | -10.2 |  |
| ZINC59207170 | -10.2 |  |
| ZINC67910942 | -10.1 |  |
| ZINC67911804 | -10.1 |  |
| ZINC67903153 | -10.0 |  |
| ZINC67910930 | -9.8 |  |
| ZINC67911581 | -9.8 |  |
| ZINC69713310 | -9.8 |  |

**Figure 5.**
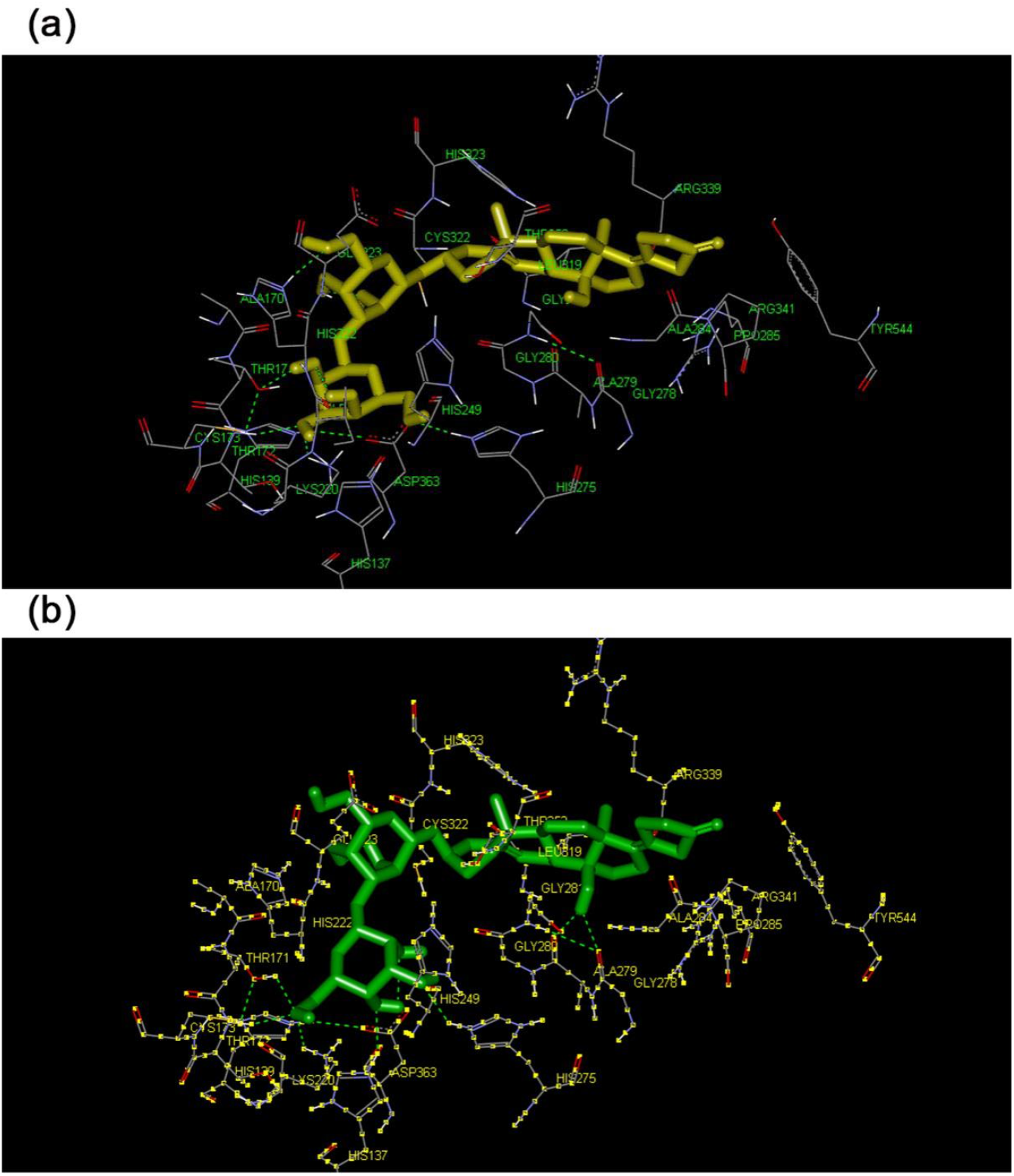
Predicted binding models of the best inhibitor ZINC67911797 (a) and its enantiomer ZINC67911804 (b) to the model structure of *Ruminococcus albus* 8 urease (RaUrease). The two inhibitors contained the same 26 amino acid residues (HIS137, HIS139, ALA170, THR171, THR172, CYS173, LYS220, HIS222, GLU223, HIS249, THR252, HIS275, GLY278, ALA284, PRO285, LEU319, CYS322, HIS323, ARG339, ARG341, ASP363, and TYR544), although ZINC67911797 had one more amino acid residue (ILE221) than ZINC67911804. There were five hydrogen bonds formed between ZINC67911797 and RaUrease, and seven hydrogen bonds formed between ZINC67911804 and RaUrease.

The amino acid residues surrounding ZINC67911797 and ZINC67911804 were conserved. The two compounds contained the same 26 residues (HIS137, HIS139, ALA170, THR171, THR172, CYS173, LYS220, HIS222, GLU223, HIS249, THR252, HIS275, GLY278, ALA284, PRO285, LEU319, CYS322, HIS323, ARG339, ARG341, ASP363, and TYR544); however, the most effective inhibitor, ZINC67911797, had one more amino acid residue (ILE221) than ZINC67911804. There were five hydrogen bonds formed between ZINC67911797 and RaUrease, and seven hydrogen bounds formed between ZINC67911804 and RaUrease. These results suggest that van der Waals forces play a more important role than hydrogen bonds in the urease-binding site. Compared with the binding model of AHA, differences in the binding site and major intermolecular forces were found to affect the efficacy of the inhibitors.

## 3. Experimental Section

### 3.1 Homology modeling

The amino acid sequence of the target protein, RaUrease, was retrieved from the NCBI database (GenBank: EGC01992.1) in FASTA format. According to the results of the BLASTp algorithm, the template protein selected was the *Helicobacter pylori* urease (HpUrease) (PDB ID: 1E9Z). The sequence alignment was performed using Clustal X 2.0 software [15], and the 3D homology model structure of RaUrease was generated using Modeller 9.16 [16].

### 3.2 Molecular dynamics simulation

Molecular dynamics simulation was performed using Gromacs 5.1 with AMBER-03 all-atom force field [17]. The temperature was maintained at 300 K. The protein was solvated using a box of simple point charge (SPC) water molecules extending at least 10 Å away from the boundary of any protein atom. An integration step of 2 fs was used. Non-bonded interactions were calculated using a cutoff of 8 Å. Long-range electrostatic interactions were calculated by Particle–Mesh Ewald summation with grid spacing of 1.2 Å and cubic interpolation. After 1000 steps of steepest descent energy minimization, the solvent and ions were equilibrated by 0.5 ns molecular dynamics simulation with the protein heavy atoms subjected to harmonic constraints under a force constant of k = 1000 kcal·mol^−1^·nm^−2^. Finally, the production run was performed for 20 ns, and the coordinates of all the atoms were stored every 100 picoseconds for further analysis.

The model quality was assessed based on the geometric quality of the backbone conformation, residue interactions, residue contacts, and energy profile of the structure using different methods, including ERRAT [18], Verify 3D [19,20], and Molprobity [21]. The volume of the binding pocket was computed using the software of the CASTp server [22] with default settings.

### 3.3 Molecular docking

AutoDock Vina was used to perform the molecular docking analyses [23]. The model structures of RaUrease and its known inhibitor AHA were determined first. The molecules were prepared as docking input files using the AutoDock Tools 1.5.6 [24]. The amino acids surrounding AHA in the binding site of RaUrease were predicted. After that, a grid box with the dimensions 126 Å × 126 Å × 126 Å was built to cover the entire model structure of RaUrease. The grid center was set at 128.255 Å × 127.264 Å × 91.45 Å to cover each active site, and a grid spacing of 0.375 Å (approximately one-fourth of the length of the carbon–carbon covalent bond) was used for the calculation of the energy map.

### 3.4 Virtual screening

The 3D structure of the established model RaUrease was used as the target protein. The Natural Products Database of the ZINC database was used because of its open source and commercial availability [25]. All compounds (11247 compounds in total) were analyzed by removing all the inorganic counter ions, adding hydrogen atoms, deprotonating strong acids, protonating strong bases, and generating stereoisomers. All compounds were then subjected to the same docking procedure. Compounds with docking scores of <-9.8 kcal mol^-1^ were selected for further virtual screening.

## 4. Conclusions

In conclusion, a reliable and reasonable 3D structure of RaUrease was successfully built with homology modeling techniques and molecular dynamics simulation methods. Twelve commercially available compounds were identified as potential urease inhibitors, and the binding site of the most effective compound on RaUrease was characterized. The findings of the present study will be of value to facilitate further experimental verification and to reduce nitrogen pollution derived from ruminant production. In addition, the present findings should help the design of novel inhibitors for enteric bacterial urease.

## Acknowledgments

This study was financially supported by the Agricultural Science and Technology Innovation Program (Project Number: ASTIP-ISA12).

